# Aggregation of misfolded proteins in the sperm head impairs preimplantation embryo development

**DOI:** 10.64898/2026.06.01.728900

**Authors:** Everardo Anta, Froylan Sosa, Jessica Drum, Kelsey Lockhart, Katy Stoecklein, Lindsey Fallon, Emma Keller, Alexandra Else-Keller, Karl Kerns, M. Sofia Ortega

## Abstract

Paternal contributions to embryogenesis extend beyond DNA, yet the molecular cargo carried by sperm and its impact on development remain poorly defined. Aggresomes (AGG), cytoplasmic inclusions formed by misfolded proteins, are present in mammalian gametes, but their functional consequences are unclear. Here, we show that excessive AGG content in bovine sperm head compromises preimplantation embryo development. Using image-based flow cytometry, sperm from 32 sires were classified into low-, moderate-, and high-AGG groups. AGG levels were unrelated to sire age and did not affect in vitro capacitation or acrosome remodeling. However, embryos derived from high-AGG sires exhibited reduced blastocyst formation, delayed cleavage timing, and a higher incidence of developmental arrest at the 4-6 cell stage. Embryos from high-AGG sires also accumulated more AGG during development, showed elevated reactive oxygen species (ROS) levels, and displayed altered mitophagy dynamics. Supplementation with an ER stress inhibitor temporarily improved cleavage but did not enhance overall blastocyst formation, indicating a limited and stage-specific effect. In vivo, embryos from high-AGG sires showed lower transferable quality compared with those from low-AGG sires. These findings establish sperm head AGG content as a novel paternal determinant of embryo quality. By linking sperm-borne misfolded protein aggregates to disrupted developmental pathways in the resulting embryo, our study reveals a previously unrecognized mechanism of paternal influence on fertility and suggests new opportunities for molecular screening for male fertility.

**Significance Statement:** Sperm contribute more to the embryo than DNA alone, yet the consequences of sperm-borne molecular cargo for early development remain largely unknown. We show that aggregates of misfolded proteins in the sperm head, a marker of disrupted protein quality control, impair preimplantation embryo development in cattle. Sires with elevated sperm aggregate content produce embryos that cleave later, arrest more frequently, and reach the blastocyst stage at lower rates, both in vitro and in vivo. These embryos carry greater aggregate loads, show heightened oxidative stress, and display dysregulated mitochondrial clearance. Our findings establish paternal proteostasis as a determinant of embryo quality and identify a class of sperm defects invisible to conventional semen analysis, opening new avenues for molecular fertility screening.

## Introduction

How parental proteostatic state influences early embryonic development remains largely unknown. Beyond fertilization, sperm deliver proteins, RNAs, and organelles that can influence developmental competence (1–5). For example, transcriptional abundance of CRISP2 and PEBP1, and low abundance of CCT8, in sperm have been positively correlated with sire fertility (6). In humans, sperm deliver a set of proteins essential for embryonic development, with dysfunction in these molecules linked to embryonic lethality (7). Among them, the transmembrane glycoprotein desmocollin 3 (DSC3) regulates the first cleavage division, while in mice, the sperm-borne microRNA-34c is also required for this process (8). These findings underscore that paternal contributions extend well beyond DNA, and emphasize the need to identify additional sperm-borne factors that may regulate early embryo development. One potential paternal factor of interest is the accumulation of protein aggregates (AGG). These AGG represent a cellular stress response rather than a single molecule. They form when misfolded proteins are actively transported to the perinuclear region via dynein and sequestered to reduce cytotoxicity (9, 10). In somatic cells, this compartmentalization is protective; however, excessive AGG accumulation can overwhelm proteostasis, disrupt Golgi and microtubule organization, and trigger reactive oxygen species (ROS) production and endoplasmic reticulum (ER) stress, ultimately impairing cellular function (11–13). When proteasomal degradation is insufficient to clear these aggregates, autophagy is activated to degrade accumulated AGG (10). In mammalian Flp-In T-REx293 cells, prolonged AGG accumulation has been linked to DNA damage, cell cycle arrest, and apoptosis (12), while in oocytes, AGG formation correlates with age-dependent proteostasis defects involving the ubiquitin-conjugating enzyme UBE2VI (14). These findings suggest that AGG formation is tightly linked to the ubiquitin-proteasome system, a pathway that is also essential for proper degradation of maternal proteins during early embryogenesis (15). Importantly, misfolded proteins and AGG have been detected in both sperm (16, 17) and oocytes (18), raising the possibility that sperm-associated protein aggregates or proteostasis defects promote AGG accumulation in the zygote, either through direct carryover or by inducing de novo aggregation during early development, and interfere with developmental events. Recent findings from our group suggest such a connection: sires with AGG accumulation in the sperm head exhibited reduced embryo production capacity. This phenotype is not detectable by routine sperm morphology and cannot be removed using standard gradient purification methods commonly employed in in vitro embryo production (19). Thus, AGG may represent a novel paternal phenotype influencing embryogenesis. The objective of this study was to determine whether sperm-borne AGG content affects early bovine embryonic development, with a particular focus on stress responses and developmental competence. It was hypothesized that high AGG content in sperm reflects underlying proteostasis defects that promote AGG accumulation and cellular stress in the developing embryo, overwhelming proteostasis, inducing ER stress, and ultimately impairing early embryonic development.

## Results

### Divergent AGG content in sires and its relationship with age

Sperm from a total of 32 Holstein sires was analyzed for AGG content using image-based flow cytometry (IBFC) and classified into three categories based on their proportion of cells positive for AGG (low-, moderate-, and high-AGG). Sires exhibited considerable variability in sperm AGG content, with high AGG content in the head ranging from 7% to 45% across the cohort of sires studied (Fig. 1A). Based on the proportion of sperm having high AGG in the head, sires were classified according to quartile classification. Sires with 7-18.1% of their sperm population positive for AGG were classified as low-AGG sires (quartile 1), whereas sires with 21-45% were classified as high-AGG sires (quartile 4). Sires falling within quartiles 2 and 3 were classified as moderate-AGG sires. To further explore whether the AGG in sperm was an age-dependent phenotype, a linear regression analysis was performed using data from 60 sires with known AGG phenotypes. The ages of the sires ranged from 11.9 to 101.9 months, and no significant correlation was found between AGG phenotype and the age of the sire (P = 0.41; Fig. 1B). This first exploratory analysis of the AGG phenotype facilitated the selection of eight sires split evenly across the two phenotypic groups (high- and low-AGG sires) for the rest of the experiments (Fig. 1C).

**Figure 1.**
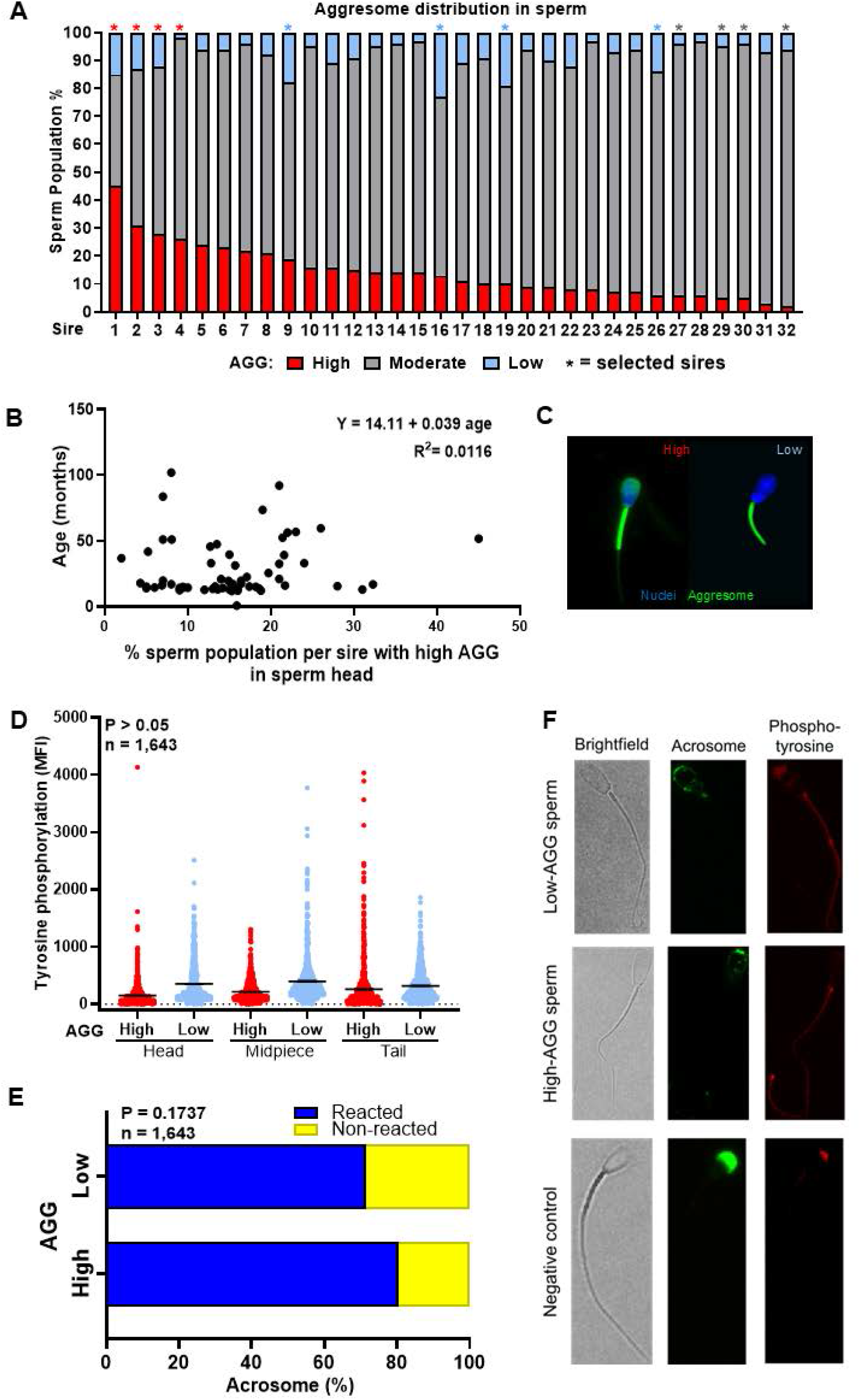
Divergent aggresome (AGG) content in bovine sperm and lack of association with age or functional competence. **(A)** Sperm from 32 sires were analyzed for whole sperm cell AGG content using image-based flow cytometry (IBFC) and classified into three categories based on the percentage of their population containing AGG: high (red), moderate (gray), and low (blue). Each bar represents the percentage distribution of sperm populations from an individual sire. Asterisks (*) denote sires selected for subsequent embryo production experiments. Sires 1-4 were selected to represent high AGG phenotype, while sires 9, 16, 20, and 26 were selected for low AGG, and sires 27, 29, 30 and 32 were used for moderate AGG. **(B)** Linear regression analysis of the age of sires and high AGG content in the sperm head. Each dot represents an individual sire. **(C)** Representative picture of a high AGG sperm and a low AGG sperm, stained with Proteostat (green) to label protein aggregates and Hoechst 33342 (blue) to label nuclear DNA, with merged images shown for visualization. **(D)** Quantification of capacitation based on tyrosine phosphorylation in sperm head, midpiece, or tail. **(E)** Quantification of acrosome-reacted and non-reacted sperm from high and low-AGG sires. **(F)** Representative images of sperm subjected to capacitation and acrosome reaction test, assessed by phosphotyrosine immunostaining (red) and peanut agglutinin (PNA; green) labeling, respectively.

### AGG does not affect sperm capacitation or acrosome membrane integrity

Next, it was evaluated whether the AGG phenotype influences functional and structural determinants of sperm fertilizing ability (Fig. 1D-F). No differences were observed in the intensity of tyrosine phosphorylation in the sperm head, midpiece, or tail between low- and high-AGG sperm. Similarly, the proportion of sperm exhibiting compromised or remodeled acrosomal membrane integrity, including patterns consistent with acrosome exocytosis, as assessed by peanut agglutinin (PNA) labeling, was not affected by AGG content.

### Effect of AGG content in sperm on embryo development

The proportion of zygotes undergoing cleavage was not affected by AGG content in sperm (P = 0.15; Fig. 2A). The ability of putative zygotes to develop into blastocysts was affected by AGG content (P = 0.0243). Further analysis showed that embryos from high-AGG sires exhibited a lower blastocyst rate compared to those from low and moderate AGG sires. Similarly, AGG content influenced the ability of cleaved embryos to develop to the blastocyst stage (P = 0.0240), with embryos from high-AGG sires showing the lowest blastocyst rate compared to those from low- and moderate-AGG sires (Fig. 2B).

**Figure 2.**
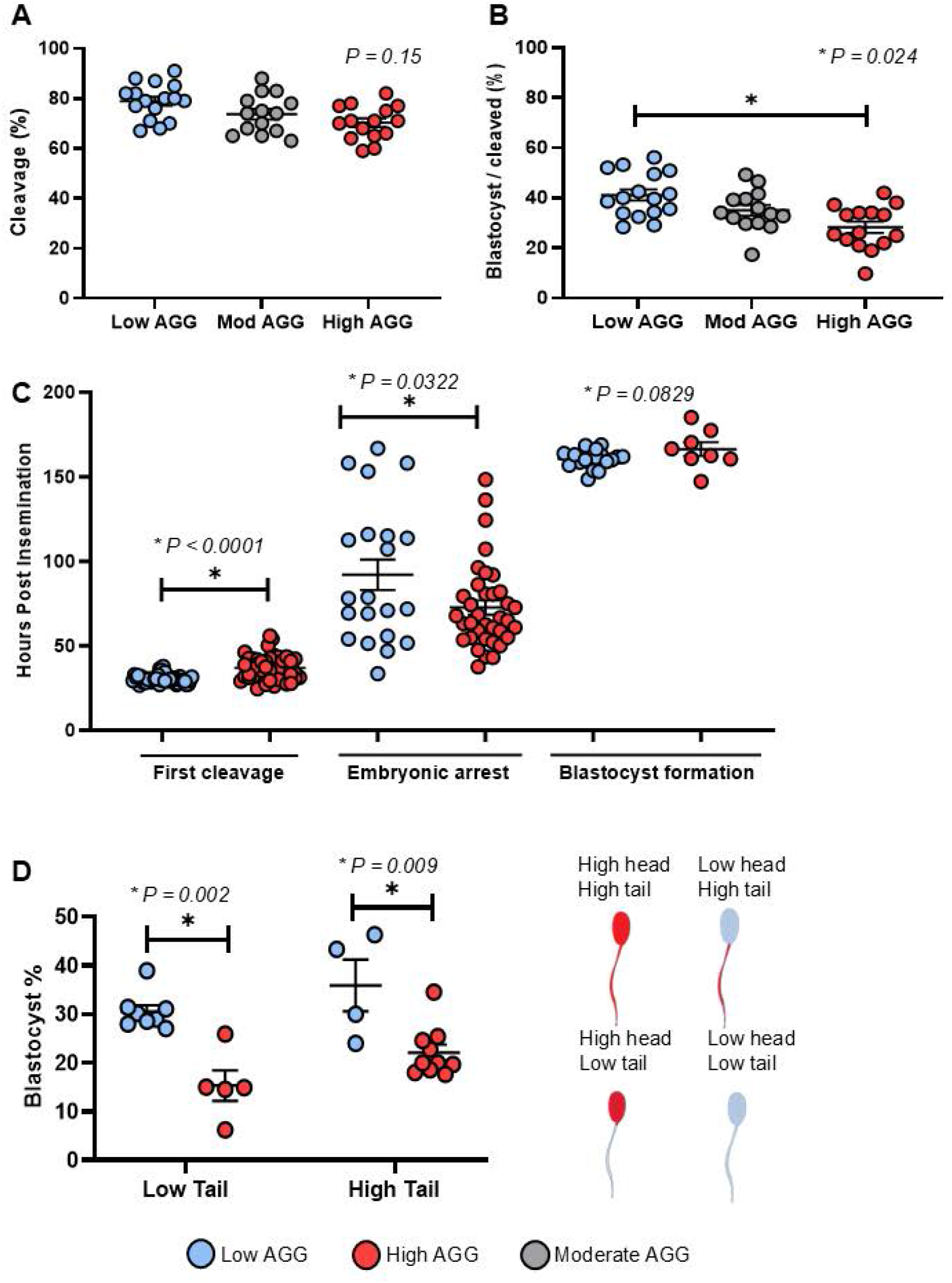
Embryo developmental competence after fertilization with distinct AGG phenotypes. **(A)** Cleavage rates of embryos generated from 12 sires classified as low (blue), moderate (gray), or high (red) based on whole sperm cell AGG content. **(B)** Percentage of embryos reaching the blastocyst stage was significantly reduced in the high AGG group (*P = 0.024). **(C)** Time-lapse records showed a significant delay in time to first cleavage and earlier embryonic arrest of embryos derived from high-AGG sires (*P < 0.0001). **(D)** Blastocyst yield of embryos produced by high and low-AGG sires stratified specifically by AGG localization within the sperm tail, independent of head AGG content.

To further examine and validate the impact of the sperm AGG phenotype throughout embryonic development, live imaging was conducted on embryos derived from high- and low-AGG sires to assess their developmental kinetics. The level of AGG content significantly delayed the timing of the first mitotic division (P < 0.0001), with embryos derived from high-AGG sires cleaving at 36.9 ± 0.8 h post insemination (hpi), compared to 30.5 ± 0.8 hpi for those from low-AGG sires (Fig. 2C). Likewise, AGG influenced the timing of embryonic arrest (P = 0.0322; Fig. 2C), with embryos derived from high-AGG sires arresting at 72.6 ± 5.5 hpi, compared to 92.02 ± 6.9 hpi for embryos from low-AGG sires. There was a tendency for AGG to affect the time of blastocyst formation (P = 0.0829) (Fig. 2C; Supplemental Material, Movie S1).

The developmental kinetics of embryos were affected by AGG, with embryos from high-AGG sires displaying a higher incidence of developmental arrest at the 4-6 cell stage than embryos from low-AGG sires (59.09% vs. 12.50%; P < 0.0003). Similarly, as previously shown, AGG affected embryonic developmental capacity (P < 0.001), with blastocyst rates lower in embryos derived from high-AGG sires compared to those from low-AGG sires. An additional question that emerged during embryo development evaluation was whether the effects of AGG were driven by overall accumulation or specific localization within the sperm (Fig. 2D). To further explore this hypothesis, six sires were evaluated based on AGG content in both head and midpiece/tail. Our findings demonstrated that elevated AGG content, specifically in the sperm head, significantly impaired embryo development regardless of midpiece/tail AGG content, as reflected in reduced blastocyst rates (P = 0.0026).

### Sperm influence on AGG accumulation in early embryonic stages

The persistence of AGG in embryos throughout development was evaluated. Embryos derived from high-AGG sires exhibited significantly higher AGG content across all developmental stages compared to embryos from low-AGG sires (P = 0.001; Fig. 3A). AGG levels peaked at the 4-6 cell stage, with an MFI nearly double that of embryos from low-AGG sires. Embryos from low-AGG sires displayed relatively stable and lower AGG levels throughout development. To further assess the contribution of sperm AGG to embryonic AGG content, biparental embryos were compared with parthenogenetic embryos at the 4-6 cell stage. The type of embryo influenced AGG accumulation (P < 0.001), with parthenogenetic embryos exhibiting lower AGG levels compared to embryos derived from either high- or low-AGG sires (Fig. 3B,C). As expected, embryos derived from high-AGG sires accumulated more AGG than those from low-AGG sires (P = 0.0001; Fig. 3B, C).

**Figure 3.**
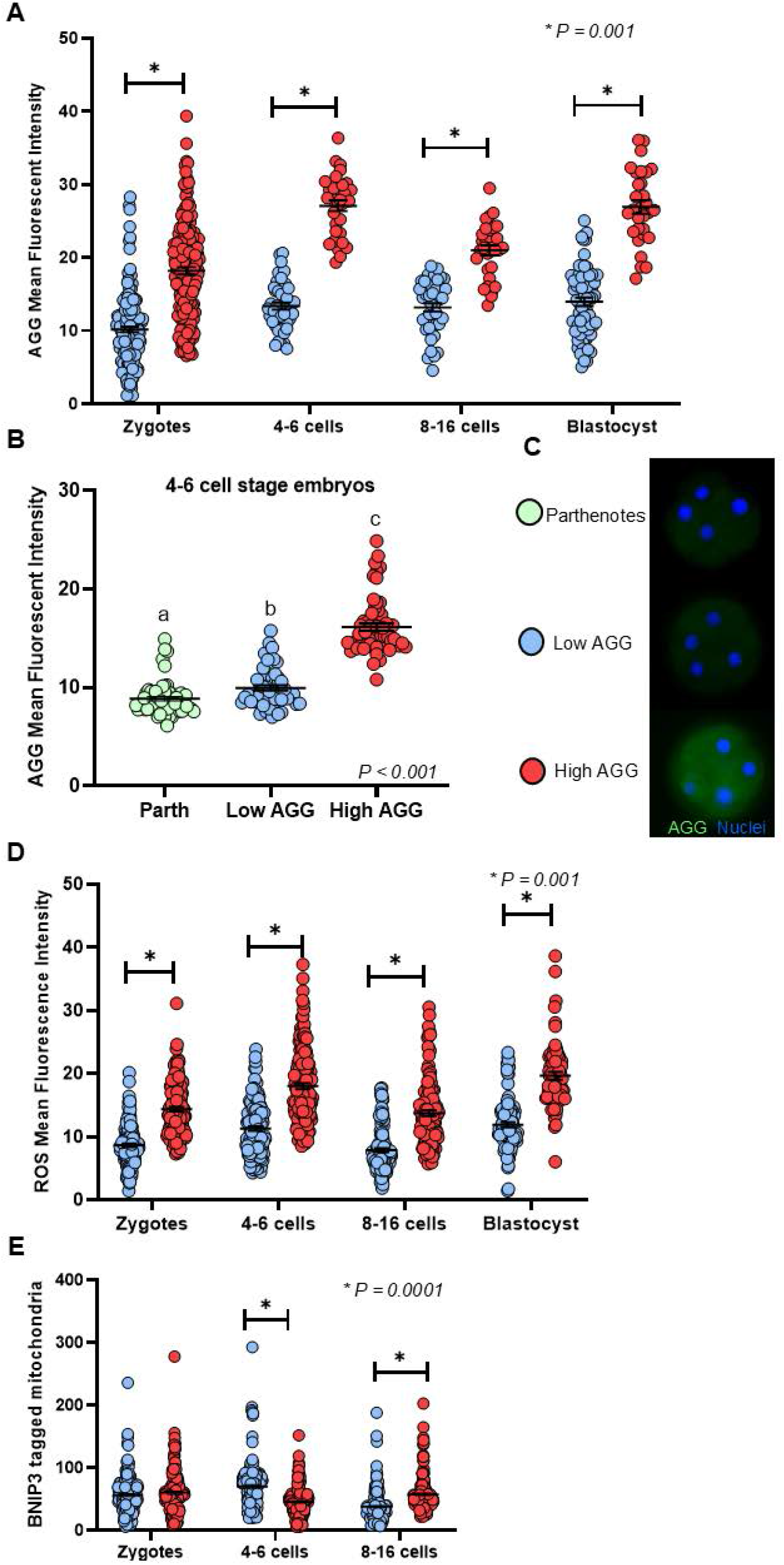
Embryos derived from high-AGG sires exhibit increased AGG content, mitochondrial stress, and ROS accumulation. **(A)** Mean AGG fluorescence intensity in embryos derived from low AGG (blue) or high AGG (red) sires, measured at the zygote, 4-6 cell, 8-16 cell, and blastocyst stages. **(B)** Mean AGG fluorescent intensity in 4-6 cell stage embryos derived from parthenotes (green), low AGG (blue), or high AGG (red) sperm. Groups with different letters (a, b, c) are significantly different (*P < 0*.*001*). **(C)** Representative images of 4-6 cell embryos stained for AGG (green) and nuclei (blue) from parthenotes, low AGG, and high AGG groups. **(D)** Reactive oxygen species (ROS) fluorescence intensity in embryos derived from low AGG (blue) or high AGG (red) sires, at the indicated developmental stages. **(E)** BNIP3-labeled mitochondria fluorescence intensity in embryos derived from low AGG (blue) or high AGG (red) sires, at the indicated developmental stages.

### Effect of AGG accumulation on oxidative stress throughout early embryo development

To evaluate the potential cytotoxic effects of AGG across different stages of embryo development, embryos at four developmental stages (zygote, 4-cell, 8-cell, and blastocyst) produced by either high- or low-AGG sires were subjected to a ROS assay. At all stages, embryos from high-AGG sires consistently exhibited elevated ROS levels compared to those from low-AGG sires (P = 0.001; Fig. 3D).

### Effects of sperm AGG on the regulation of mitophagy activity across early embryo development

Mitophagy activity was not affected by sperm AGG (P = 0.9015). Stage of development influenced the mitophagy activity (P = 0.0001); however, a significant interaction between sperm AGG and stage of development was observed (P = 0.0001). Further analysis showed that embryos derived from low-AGG sires displayed a peak in mitophagy activity at the 4-6 cell stage, followed by a significant reduction at the 8-16 cell stage, consistent with the expected temporal dynamics of mitochondrial clearance. In contrast, embryos from high-AGG sires exhibited a delayed mitophagy activity, with a significant mitophagy activity increase at the 8-16 cell stage compared to the embryos from low-AGG group (P = 0.0001; Fig. 3E).

### ER stress inhibition partially improves cleavage rate but does not alter autophagy or ER stress markers

To determine whether alleviating ER stress could improve embryonic development in embryos derived from high-AGG sires, culture medium was supplemented with an ER stress inhibitor (Tauroursodeoxycholic acid-TUDCA). Inhibiting ER stress affected cleavage rates (P = 0.0143). TUDCA supplementation increased cleavage rates in embryos from high-AGG sires, which exceeded those of both untreated high-AGG controls and TUDCA-treated embryos from low-AGG sires (Fig. S1). Treatment with TUDCA did not affect the ability of putative zygotes or cleaved embryos to reach the blastocyst stage (P > 0.05). Furthermore, TUDCA treatment did not affect the gene expression of key autophagy and ER stress-related markers (P > 0.05) (Fig. S1).

### High AGG in the sperm head affects in vivo embryo production

To determine whether sire AGG content influences the developmental quality of embryos produced in vivo, the proportion of transferable versus non-transferable embryos was evaluated. Sires with high AGG content produced embryos of lower developmental quality, as reflected by an increased proportion of non-transferable embryos and a corresponding reduction in transferable embryos relative to low-AGG sires (Fig. 4).

**Figure 4.**
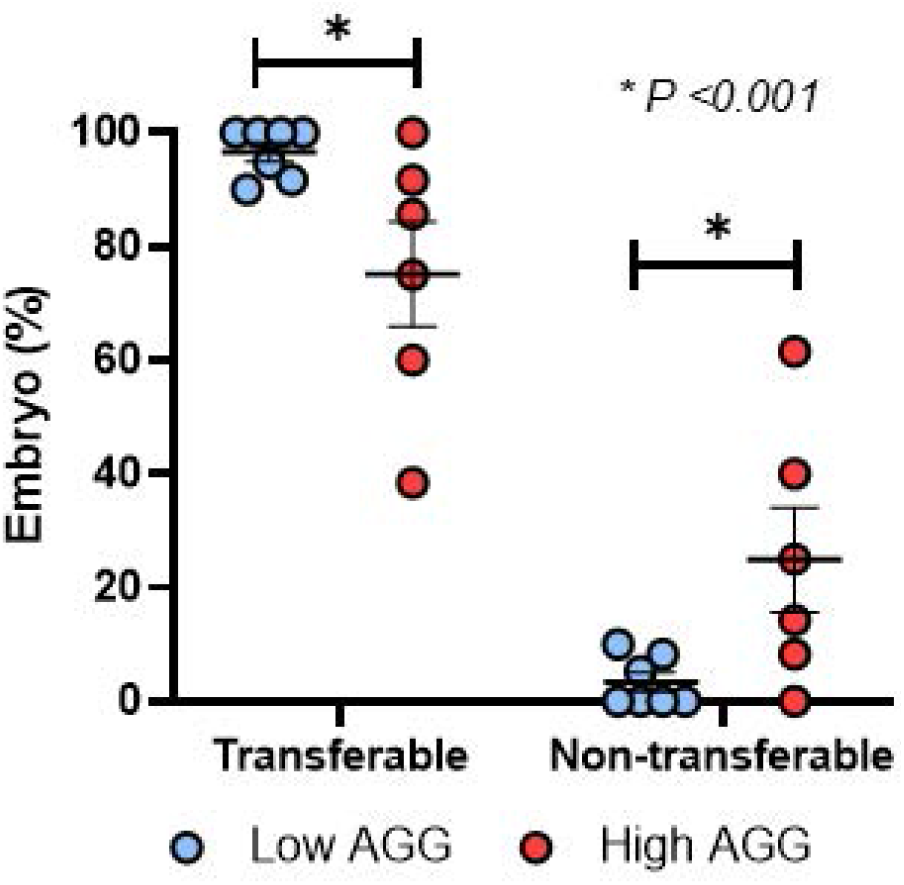
Embryos derived from high-AGG sires exhibit reduced in vivo developmental competence. Percentage of transferable and non^+^transferable embryos recovered after embryo transfer, derived from low AGG (blue) or high AGG (red) sperm.

## Discussion

The objective of this study was to elucidate the effects of AGG content in the sperm head on embryo development. Formation of AGG is a cellular protective mechanism that arises in response to proteostasis disruption. However, sperm can deliver AGG into the oocyte, affecting subsequent embryo development. A recent study in mice reported that AGG are effectively cleared from oocytes following maturation and before the first mitotic division (18). Thus, the persistent presence of AGG observed in bovine embryos derived from high-AGG sires in the present study reveals that sperm-borne AGG can be transferred into the oocyte, and that there may be a failure in AGG clearance mechanisms during early embryo development. In mice, the failure to clear protein aggregates in autophagy-deficient embryos results in developmental arrest at the 2-cell stage (18). In the present study, a high proportion of embryos arrested at the 4-6 cell stage, suggesting that bovine embryos may have an impaired autophagy mechanism for AGG clearance, although an active clearance mechanism analogous to that described in mice cannot be ruled out.

The presence of AGG in the oocyte suggests that early accumulation may initiate a progressive build-up during subsequent embryo development. Further research is required to elucidate the interaction between oocyte- and sperm-derived AGG and their consequences for embryo development.

It has been shown that protein aggregates, including AGG, can partition during mitotic divisions and persist across cell lineages (20). This inheritance of AGG may amplify cellular stress on the proteostasis pathways of daughter cells, particularly when the autophagic machinery is overburdened or compromised, potentially promoting a build-up phenomenon as proposed earlier. It is well accepted that mitophagy is completed by the 4-6 cell stage of embryonic development (21). An important consideration is whether sperm AGG are directly responsible for the observed embryonic phenotypes or instead reflect broader defects in sperm quality. While our findings show a clear association, they do not distinguish between direct transfer of AGG into the oocyte and de novo AGG formation induced by sperm-derived factors. The comparison with parthenogenetic embryos supports a paternal contribution, but it is not definitive, as parthenotes differ from biparental embryos in key aspects such as centriole inheritance and cell cycle regulation, which may independently affect AGG levels. Thus, increased AGG in biparental embryos is consistent with, but does not prove, physical transmission. Future studies using approaches such as fluorescent labeling and tracking of sperm-derived AGG will be required to directly test this mechanism.

In the present study, the mitophagy level in the high-AGG group was significantly lower at the 4-6 cell stage of development but increased at the 8-16 cell stage. This suggests that an efficient autophagy response may be crucial during the early stages of embryo development to support normal progression. Although mitophagy was elevated later, at the 8-16 cell stage, this delayed response may be insufficient to restore proteostasis, ultimately compromising embryo development. This finding may also indicate compromised autophagic machinery resulting from AGG accumulation.

Our study shows that AGG may also trigger additional forms of cellular stress, as elevated levels of ROS were observed in embryos derived from high-AGG sires throughout early development. A recent study from our laboratory reported that embryos from high-AGG sires exposed to short-term heat shock exhibited increased ROS levels compared to heat-shocked embryos from low-AGG sires (22). These findings suggest that embryos derived from high-AGG sires may possess intrinsic vulnerabilities that predispose them to heightened cellular stress responses even to minimal external stimuli, thereby disrupting homeostatic balance and leading to impaired embryo development, as observed in the present study. Mechanistically, misfolded proteins entering the ER can further amplify ROS generation through mediators such as protein disulfide isomerase (PDI) and NADPH oxidase 4 (Nox4) (23).

This ER stress, combined with inefficient degradation of misfolded proteins, may establish a feedback loop that amplifies oxidative damage, exacerbates cellular dysfunction, and further impairs development. The interplay between ER stress and ROS production, as described by Zeeshan et al. (23), aligns with the elevated oxidative stress observed in embryos from high-AGG sires. These data support the idea that AGG accumulation might compromise autophagic pathways and overwhelm ER protein-folding capacity, contributing to developmental arrest and reduced blastocyst yield.

Although embryos from high-AGG sires exhibited elevated ROS levels and impaired development, we did not observe significant differences in the expression of ER stress and autophagy markers between groups. This discrepancy could reflect a temporal disconnect between the onset of oxidative stress and the transcriptional or translational activation of ER stress and autophagy responses. For instance, these pathways may be transiently activated at earlier time points and return to baseline by the time of sampling, especially in fast-developing systems like early embryos.

Another possibility is that embryos from high-AGG sires fail to display an adequate adaptive response to stress. In this case, although stress is present, the expected upregulation of protective mechanisms like the unfolded protein response or autophagy does not occur effectively, which could explain both the unchanged marker levels and the increased susceptibility to developmental arrest. In addition, these findings may also indicate that AGG clearance in embryos relies on mechanisms outside the canonical ER stress response. Alternative pathways, such as chaperone-mediated mechanisms or non-canonical proteasomal degradation, might play a role in mitigating AGG accumulation. It also raises the possibility that the gene panel assessed does not fully represent the complexity of these responses. Importantly, these results indicate that the molecular mechanisms linking AGG accumulation to impaired embryo development remain incompletely resolved. Future studies incorporating expanded transcriptomic or proteomic analyses across multiple developmental time points will be necessary to determine whether ER stress and autophagy pathways are directly involved or whether alternative mechanisms underlie the observed phenotypes.

Interestingly, supplementation with the ER stress inhibitor TUDCA improved cleavage rates in embryos derived from high-AGG sires but did not enhance blastocyst formation or alter expression of ER stress and autophagy markers. These findings suggest a potential, stage-specific contribution of ER stress to early cleavage events; however, this experiment was not designed to rigorously test a rescue mechanism. The use of a single dose and timing limits interpretation, and therefore, these results should be considered preliminary. Additional studies exploring different treatment windows and concentrations will be required to determine whether modulation of ER stress can meaningfully improve developmental outcomes.

The increased incidence of developmental arrest at the 4-6 cell stage in embryos derived from high-AGG sires suggests that paternal AGG contribution triggers a response in the embryo that ends up disrupting development at early stages. Consistent with these findings, our in vivo embryo production data revealed a significantly lower proportion of transferable embryos derived from high-AGG sires compared to low-AGG sires, indicating that the negative impact of AGG extends beyond in vitro conditions. The stage of cellular arrest in these embryos coincides with the minor genome activation (MGA) and precedes the major embryonic genome activation (EGA), a pivotal period when transcriptional control shifts from maternal transcripts to embryonic gene products. During MGA, the degradation of maternal mRNAs and chromatin remodeling creates a transcriptionally permissive environment necessary for EGA (24, 25).

Elevated AGG content may disrupt these processes, either by interfering with proteostatic mechanisms required for maternal mRNA clearance or by altering chromatin accessibility, delaying the activation of key embryonic genes. The consistent timing of blastocyst formation across groups further suggests that AGG’s adverse effects are most pronounced during the pre-EGA stage, with developmental arrest occurring before the embryo can transition to autonomous transcription. While sperm are traditionally viewed as a mere vehicle for genetic material, this study highlights how sperm could regulate the embryo’s transcriptional landscape.

In conclusion, the present study highlights the significant impact of elevated AGG content in sperm on early embryonic development, with embryos derived from high-AGG sires exhibiting reduced developmental competence and increased susceptibility to arrest. The inability of embryos to effectively manage paternal AGG load may contribute to arrest at the 4-6 cell stage before EGA. Although we cannot confirm that all embryos fertilized with high-AGG sires exclusively inherited elevated AGG content, our results suggest that when as little as 21% of the sperm population is affected by high AGG levels, there is a clear reduction in development to the blastocyst stage (Fig. S2). Furthermore, increased AGG predisposes embryos to proteostasis defects that trigger cellular stress pathways, including increased oxidative stress and delayed mitophagy. Further research is needed to isolate AGG-rich sperm populations to evaluate their developmental competency directly. Such studies will help elucidate the mechanistic role of AGG in driving early embryonic arrest and provide insights into potential interventions to mitigate its effects. Finally, the origin of AGG accumulation in the sperm head remains unclear, as does whether it can be managed to improve embryonic development in vitro and in vivo.

## Materials and Methods

All reagents were obtained from Sigma-Aldrich (MO, USA) unless otherwise stated. All statistical analyses were performed using the Statistical Analysis System SAS V 9.4 (SAS Institute, NC, USA). Significance was considered when P < 0.05. Results are presented as least-squares means ± standard error of the mean (SEM).

### Sire phenotyping and AGG labeling for sample acquisition

Frozen semen straws from thirty-two commercial Holstein Friesian sires were donated from Select Sires Inc. (OH, USA). Sire age ranged from 12.5 to 73.5 months at the time of collection, and they were evaluated for AGG content using IBFC. A total of two straws, with 10,000 sperm cells, were evaluated per sire.

To determine AGG content in sperm, semen straws were thawed in a 38.5°C water bath and transferred into 15 mL tubes containing 7 mL of pre-warmed phosphate-buffered saline (PBS) (Thermo Fisher Scientific, MA, USA). Samples were centrifuged at 2,200 rpm (541 g) for 7 min using a swinging-bucket rotor. After centrifugation, the supernatant was discarded, and the pellet was resuspended in 7 mL PBS. This washing step was repeated under the same conditions. Immediately after, sperm were resuspended in 7 mL of 2% (v/v) formaldehyde solution and fixed at room temperature (RT) for 40 min. After fixation, samples were centrifuged as mentioned above, and supernatant discarded. The pellet was washed in 7 mL of PBS containing 0.05% (w/v) sodium azide (NaN_3_), centrifuged again, resuspended, and stored in 7 mL of PBS with NaN_3_ at 4°C until analysis.

To detect AGG in fixed sperm, samples were stained with the PROTEOSTAT aggresome detection kit (Enzo Life Sciences, NY, USA) at a dilution of 1:2000, and DNA was stained with 1 µg/mL of Hoechst 33342 (H33342; Invitrogen Thermofisher Scientific). IBFC was performed using the ImageStream®X MkII image-based flow cytometer (ISX; Fremont, CA, USA), equipped with a 12-channel detection system and a 40x objective (numerical objective of 0.9), with an image rate up to 2,000 events per sec. The sheath fluid used was PBS without Ca^2+^or Mg^2+^. The flow-core diameter and speed were 10 μm and 66 mm per sec., respectively. The following laser settings were used: 405 nm laser at 40 mW to excite H33342, 488 nm laser at 50 mW to excite the PROTEOSTAT AGG probe, and 785 nM side scatter laser at 10 mW. Raw image data were acquired using INSPIRE® software. To produce the highest resolution, the camera setting was at 0.5 μm per pixel of the charged-coupled device. In INSPIRE® ISX data acquisition software, two brightfield channels were collected (channels 1 & 9), one AGG image (channel 2), one side scatter (SSC; channel 6), and one H33342 (channel 7), with a minimum of 10,000 spermatozoa collected.

Data were analyzed using AMNIS IDEAS® analysis software version 6.4 from Cytek (Fremont, CA, USA). An initial gating step excluded events that were not single, in-focus sperm cells, as well as those that were laterally aligned with the camera, from downstream analysis. For image-based analysis, custom-defined masks were applied to distinguish the head and tail of each sperm cell. Head-only masks were defined by taking the mask created by nuclear stain H33342, adding a 2-pixel dilation, and then analyzing the image values of other fluorescent/brightfield images. Likewise, a mask for the tail only was created by subtracting this sperm head mask from the remainder of the sperm as described (15).

### Tyrosine phosphorylation and acrosome reaction assays

Frozen semen from eight sires (four low-AGG and four high-AGG) was thawed at 37 °C for 30 s. Sperm were purified using an isolate gradient through centrifugation at 1,000 × g for 5 min. The resulting pellet was washed twice in HEPES-TALP medium at 500 × g for 3 min, and the supernatant was removed. To promote sperm capacitation, sperm were incubated in 960 µL of IVF medium supplemented with 50 µM calcium ionophore A23187 (calcimycin, Sigma-Aldrich, Cat. No. C7522) and 40 µL of PHE solution (0.5 mM penicillamine, 0.25 mM hypotaurine, 0.25 µM epinephrine). Negative-control sperm were incubated only in IVF medium. Both groups were incubated overnight in a humidified atmosphere of 5% CO_2_ in air at 38.5 °C. For tyrosine phosphorylation assays, sperm were incubated overnight with 5 µg/mL of anti-phospho-tyrosine mouse antibody (#9411, Cell Signaling Technology), followed by labeling with goat anti-mouse IgG secondary antibody conjugated to Alexa Fluor™ 594. For acrosome reaction assays, sperm were stained with 0.5 µg/mL peanut agglutinin CF®488 (PNA; Biotium, Cat. No. 29060) and 1 µg/mL Hoechst for 30 min. After staining, sperm were washed and smeared on slides and mounted with coverslips. Sperm were imaged using a fluorescence microscope, and images were analyzed with ImageJ software. For tyrosine phosphorylation, mean fluorescence intensity in the head, midpiece, and tail was quantified. For the acrosome reaction, sperm showing fluorescence at the acrosome were classified as unreacted, whereas those lacking acrosomal fluorescence were classified as having undergone the acrosome reaction.

### Sire classification per AGG content in sperm head

Sires were classified based on the percentage of sperm in their population displaying AGG content in the sperm head. Classification was performed using quartile-based ranking with the RANK procedure of SAS (SAS Institute Inc., Cary, NC). Quartile 1 represented sires with the lowest AGG content, with 7% to 18.1% of their total sperm population exhibiting high AGG content, while Quartile 4 included sires with the highest AGG content ranging from 21% to 45%. Sires from Quartile 2 and 3 representing moderate AGG content, had over 85% of their total sperm population classified as moderate AGG. Based on their quartile classification, sires were assigned to one of three groups: high, moderate, or low AGG content. These groups were used for subsequent analysis.

### In vitro embryo production with sires with divergent AGG content

To evaluate the effects of AGG content in the sperm head on embryo development, embryos were produced with a total of 12 sires, four of each of the three AGG phenotypes across 17 rounds of in vitro embryo production (IVP). The low AGG sire group was treated as IVP control, and when failed, that round was not considered for analysis. At least 2 sires per classification used per round of IVP. In vitro bovine embryos were produced as described elsewhere (19, 26, 27). Briefly, cumulus oocyte complexes (COCs) were retrieved from abattoir-derived ovaries by slicing the ovarian surface and rinsed into oocyte-collection medium. Grade 1 and 2 COCs were selected based on the International Embryo Technology Society (IETS) guidelines, ensuring at least two cumulus layers and homogeneous cytoplasm. Selected COCs were divided into groups of 50 and cultured in 5 well dishes containing 500 μl of pre-equilibrated oocyte maturation medium overlaid with 250 μl of mineral oil (Irvine Scientific, CA, USA). Maturation took place in an incubator in a humidified atmosphere of 5% CO2 in air at 38.5°C for 22-24 hours. After maturation, COCs were divided into three groups, and each group was fertilized with a high, moderate, and low AGG sire, respectively. For all sires, the final sperm concentration in the fertilization plate was 1 × 106/ml, and sperm and COCs were coincubated for 16-18 h. Following fertilization, cumulus cells were removed and presumptive zygotes were cultured in groups of 50 in 500 μl of synthetic oviduct fluid (SOF-BEII) overlaid with 250 μl of mineral oil. Embryos were cultured for 7 days in an incubator in a humidified atmosphere of 5% CO2, 5% O2, 90% N2 at 38.5 °C. A total of 3422 presumptive zygotes were cultured during the entire experiment (low AGG n = 1235; moderate AGG n = 1081; high AGG n = 1106), cleavage rates were assessed on Day 3 (Day 0 = fertilization day), and blastocyst rate was evaluated on Day 7.5 of development.

### Effect of AGG content in sperm head on embryo kinetics using time-lapse imaging

To validate the effects of AGG content on development, embryos were produced as described above and cultured in a time-lapse incubator (EmbryoScope, Vitrolife, Denmark). Images were captured using a 10-minute cycle for up to 180 hpi. Two independent IVP rounds were conducted using a total of 144 presumptive zygotes, 72 per round (36 presumptive zygotes/sire) with one high and one low AGG sire to compare the extreme phenotypes. Developmental parameters for individual embryos evaluated were time to first cleavage, time of developmental arrest, the stage at which the arrest occurred, and time to blastocyst formation.

### Sperm AGG localization or accumulation and age regression for AGG content

To investigate whether the AGG present in sperm affected embryo development due to the localization of AGG (head or tail) or their accumulation (high or low), six sires were selected by categorizing them into four groups based on AGG content localization: high AGG in both the head and tail, high AGG in the head but low in the tail, low AGG in the head but high in the tail and low AGG in both the head and the tail. In addition, to evaluate if the high AGG phenotype observed in sires was age-dependent, a linear regression analysis was conducted using data from 60 sires phenotyped for AGG content. The ages of the sires ranged from 11.9 to 101.9 months, with all sires having prior IVP data from our group (26, 28). This dataset included the 12 sires used across the experiments presented in this study.

### Evaluation of AGG persistence across embryo development and mitochondrial clearance

To investigate AGG content persistence across different stages in early embryo development, embryos were produced using four high-AGG sires and four low-AGG sires, across three independent rounds of IVP. A total of 1,365 embryos were collected at four developmental stages: zygotes at 14 hpi (low AGG n = 165; high AGG n = 176), 4-6 cell embryos at 38 hpi (low AGG n = 169; high AGG n = 172), 8-16 cell embryos at 62 hpi (low AGG n = 183; high AGG n = 173), and blastocysts at 182 hpi (low AGG n = 195; high AGG n = 132). Embryos were fixed in 4% paraformaldehyde (PFA) for 20 min at RT. After fixation, embryos were washed in PBS-PVP, and then incubated for 1h in a fixation/permeabilization solution (FOXP3 Transcription Factor Staining Buffer Set, Invitrogen, MA) following manufacturer’s specifications. Following incubation, embryos were washed three times in 1X permeabilization buffer (Invitrogen). Subsequently, embryos were incubated in a blocking buffer (5% bovine serum albumin [BSA] in PBS) for 15 min at RT, followed by three additional washes in permeabilization buffer (Invitrogen). Additionally, to determine if embryonic arrest and AGG accumulation in the embryo were driven by impaired paternal mitochondria degradation, staining for mitophagy was performed on zygotes, 4-6 cell stage, and 8-16 cell stage embryos. Embryos were incubated with the mitophagy mediating receptor BNIP3 primary antibody (1:800) for 1 hour at RT. Embryos were washed three times in PBS-PVP and then incubated with Alexa Fluor 555 anti-rabbit IgG secondary antibody (1:500) for 1 h at RT. Embryos, including zygotes (17 hpi), 4-6 cells, 8-16 cells, and blastocysts, were incubated in PROTEOSTAT (1:2000; Enzo Life Sciences) for 30 min at RT for AGG detection. Finally, all embryos were counterstained with Hoechst (Invitrogen) for 10 min to label nuclear DNA and washed three times in PBS-PVP.

Embryos were then mounted on slides, and fluorescence imaging was performed on the same day using a Keyence microscope with the following filters: CY5 (620/60nm) for mitophagy staining, GFP (470/40nm) for AGG content, and DAPI (360/40nm) for nuclear staining. Images were analyzed using FIJI (ImageJ V. 1.54k; National Institutes of Health, Bethesda, MD) software to quantify fluorescence intensity. Mean fluorescence intensity (MFI) was calculated by subtracting background fluorescence from cytoplasmic fluorescence. For mitophagy, individual events were counted throughout 12, 0.5 μm Z-stack sections of the embryo.

### Assessment of ROS species in embryos

To evaluate the potential cytotoxic effects of AGG on embryonic development, ROS levels were quantified as markers of oxidative stress. A total of 1,042 embryos were produced across three replicates using the same sires as in the previous experiments (four low AGG and four high AGG). Embryos were live-stained at different stages of development (zygotes, low AGG n = 147, high AGG n = 144; 4-6 cells, low AGG n = 157, high AGG n = 159; 8-16 cells, low AGG n = 137, high AGG n = 122; and blastocysts, low AGG n = 105, high AGG n = 71) to measure ROS levels.

Embryos were incubated for 30 min under culture conditions in CellROX Deep Red reagent at 5 μM (ThermoFisher Scientific, Invitrogen), diluted in SOF-BEII at 38.5°C. Following ROS staining, embryos were fixed as previously described and co-stained using PROTEOSTAT (1:2000) as previously described. Images were captured within two hours of staining.

### Generation of parthenogenic embryos to evaluate paternal AGG contribution

To evaluate the paternal contribution of AGG, oocytes were recovered and matured as previously described in four independent IVP replicates. Mature oocytes were divided to be fertilized as follows: fertilized with a low AGG sire, fertilized with a high AGG sire, and parthenogenetically activated. For parthenogenic activation, matured COCs were denuded, washed in HEPES-TALP, and incubated in HEPES-TALP containing 58.55 mM sucrose (Fisher Scientific) for two min. Immediately after, activation was performed by treating the oocytes with 5 mM calcium ionomycin (Sigma-Aldrich, MO, USA) for 5 min. Following ionomycin treatment, oocytes were washed in HEPES-TALP and incubated in 1.95 mM +6-DMAP (Sigma-Aldrich) in SOF-BEII for three hours to inhibit protein synthesis. Activated oocytes were cultured in SOF-BEII under standard conditions. After 26 hours of culture, 4-6 cell-stage embryos were recovered from each group: low AGG-derived embryos (n=52), high-AGG embryos (n=60), and parthenotes (n=60). Embryos were fixed and stained to assess AGG and counterstained with Hoechst. Fluorescence intensity was subsequently analyzed to quantify AGG levels.

### ER stress inhibition as a strategy to mitigate the high AGG phenotype in embryos

To investigate whether developmental outcomes could improve by inhibiting ER stress, Tauroursodeoxycholic acid (TUDCA) (10 μM) was added to the culture medium at the beginning of culture. A total of 1,572 presumptive zygotes produced from two sires (one high and one low AGG) across three IVP procedures were assigned to one of four experimental groups: low AGG ± TUDCA, and high AGG ± TUDCA. From each round, embryos were split for two different experiments, development and gene expression. The total number of embryos used for development were: low AGG -TUDCA (n = 200); low AGG +TUDCA (n = 300); high AGG -TUDCA (n = 211); and high AGG +TUDCA (n= 280). Embryo development was monitored, and cleavage rates were assessed on Day 3 post-insemination, while blastocyst production was evaluated on Day 7.5.

### Effect of ER stress inhibitor on ER stress and autophagy pathways

To corroborate the impact of TUDCA treatment on embryos derived from high and low-AGG sires, gene expression of key ER stress and autophagy markers was analyzed at the 4-6 cell stage. For each of the three replicates, 20-25 embryos per group (low AGG -TUDCA n = 140; low AGG +TUDCA n= 146; high AGG -TUDCA n = 150; and high AGG +TUDCA n = 145), were harvested and pooled in two separate groups to ensure sufficient RNA yield for qPCR analysis. ER stress-related genes, including endoplasmic reticulum to nucleus signaling 1 (*ERN1*), eukaryotic translation initiation factor 2 alpha kinase 3 (*EIF2AK3*), activating transcription factor 6 (*ATF6*), as well as autophagy-related genes including microtubule-associated protein 1 light chain 3 beta (*MAP1LC3B*), sequestosome 1 (*SQSTM1*), and beclin 1 (*BECN1*), were assessed.

RNA was isolated from embryos using the Qiagen AllPrep DNA/RNA Mini Kit (Qiagen, Germany) as described elsewhere (29). cDNA was synthesized from RNA using the LunaScript RT SuperMix Kit (New England Biolabs, MA, USA) according to the manufacturer’s instructions. A no-reverse transcriptase (no-RT) control was prepared from each sample by substituting the LunaScript RT SuperMix with 4 μL of No-RT Control Mix (5X). cDNA synthesis was performed in a thermal cycler (Bio-Rad T100; CA, USA) using the following conditions: 25°C for 2 min, 55°C for 10 min, and 95°C for 1 min. Gene expression was determined by RT-qPCR using a CFX-96 Real-Time PCR System (Bio-Rad). Primers were designed and validated as previously described (30). The primer sequences used are shown in Table S1.

Each qPCR reaction was performed in a 10 µL total volume, containing 5 µL of SYBR Green Master Mix (GlpBio, CA, USA), 3 µL of cDNA, 0.5 µL of each primer, and 1 µL nuclease-free water (IDT, IA, USA). All reactions were run in duplicates with each treatment group represented by three biological replicates. The thermocycling conditions were as follows: initial denaturation at 95°C for 10 min, followed by 40 cycles of denaturation at 95°C for 15 seconds, and annealing/extension at 60°C for 1 min, with a melt curve analysis. Gene expression levels were normalized to the housekeeping gene, ACTB, and relative expression levels were calculated using the 2^-ΔΔCT method using the low AGG control as reference.

### In vivo validation of AGG effects on embryo development

To determine if the impact of AGG phenotype was restricted to IVP, sires previously used to produce embryos in vivo (28) were phenotyped for AGG content as described above. For in vivo embryo production, 20 heifers in a crossover design, averaging 18 months of age, were synchronized, superstimulated, inseminated, flushed, and embryos graded as described in a previous study (28). A total of 101 embryos were classified as transferable or non-transferable. A total of eight low-AGG sires and five high-AGG sires were represented in this dataset. Unfertilized oocytes (UFOs) made up 11% of the structures recovered and were evenly distributed across sires and heifers, and were removed from analysis.

### Statistical analysis

All statistical analyses were performed using SAS version 9.4 (SAS Institute Inc., Cary, NC, USA). A significance level of P < 0.05 was considered statistically significant. When a significant difference was detected, Tukey’s multiple comparison test was applied. Results are reported as least-squares means ± SEM. Cleavage rate, embryo development, AGG localization, accumulation, and effect of AGG on embryokinetics were analyzed using analysis of variance (ANOVA) with the GLIMMIX procedure in SAS. Sire’s AGG phenotype was treated as a fixed effect, while sire and IVP round were treated as random effects. A linear regression analysis was used to examine the relationship between age and high AGG phenotype. Tyrosine phosphorylation intensity in the sperm head, midpiece and tail were analyzed using a PROC MIXED procedure of SAS. This included AGG group as a fixed effect and sire nested within AGG as a random effect. Least-squares means were compared using Tukey’s adjustment for multiple comparisons. The proportion of reacted spermatozoa was analyzed using a PROC GLIMMIX procedure of SAS with a binomial distribution and logit link function, including sire as a random effect and AGG group as a fixed effect. The effects of AGG phenotype and developmental stage on embryonic AGG content and mitophagy were analyzed by ANOVA. Fixed effects included replicate, AGG phenotype, developmental stage, and their interaction. To evaluate the effect of AGG phenotype on ROS accumulation, ANOVA was conducted using the GLM procedure of SAS. Fixed effects included AGG phenotype, developmental stage, and their interaction. To determine the paternal contribution of AGG to the resulting embryos, ANOVA was performed using the GLM procedure. Fixed effects included experimental group, replicate, and their interaction. For gene expression analysis, differences among treatments were assessed using ANOVA via the GLIMMIX procedure. Fixed effects included treatment (low AGG -TUDCA, low AGG +TUDCA, high AGG -TUDCA, and high AGG +TUDCA) and replicate. For in vivo-derived embryos, the effect of AGG phenotype was evaluated using the GLIMMIX procedure, with AGG phenotype considered as a fixed effect.

## Acknowledgments

This project was supported by the Agriculture and Food Research Initiative (AFRI) Competitive Grant no. 2022-67015-36298 (K.K.) from the USDA National Institute of Food and Agriculture (NIFA), Hatch WIS05090 (E.A.), and the Dairy Innovation Hub (F.S.). We thank Lindsey Jennett for her assistance with image-based flow cytometry data acquisition and data analysis.

## Data Availability Statement

All data are contained within the manuscript and supplemental materials.

## Figure Legends

**Figure S1.**
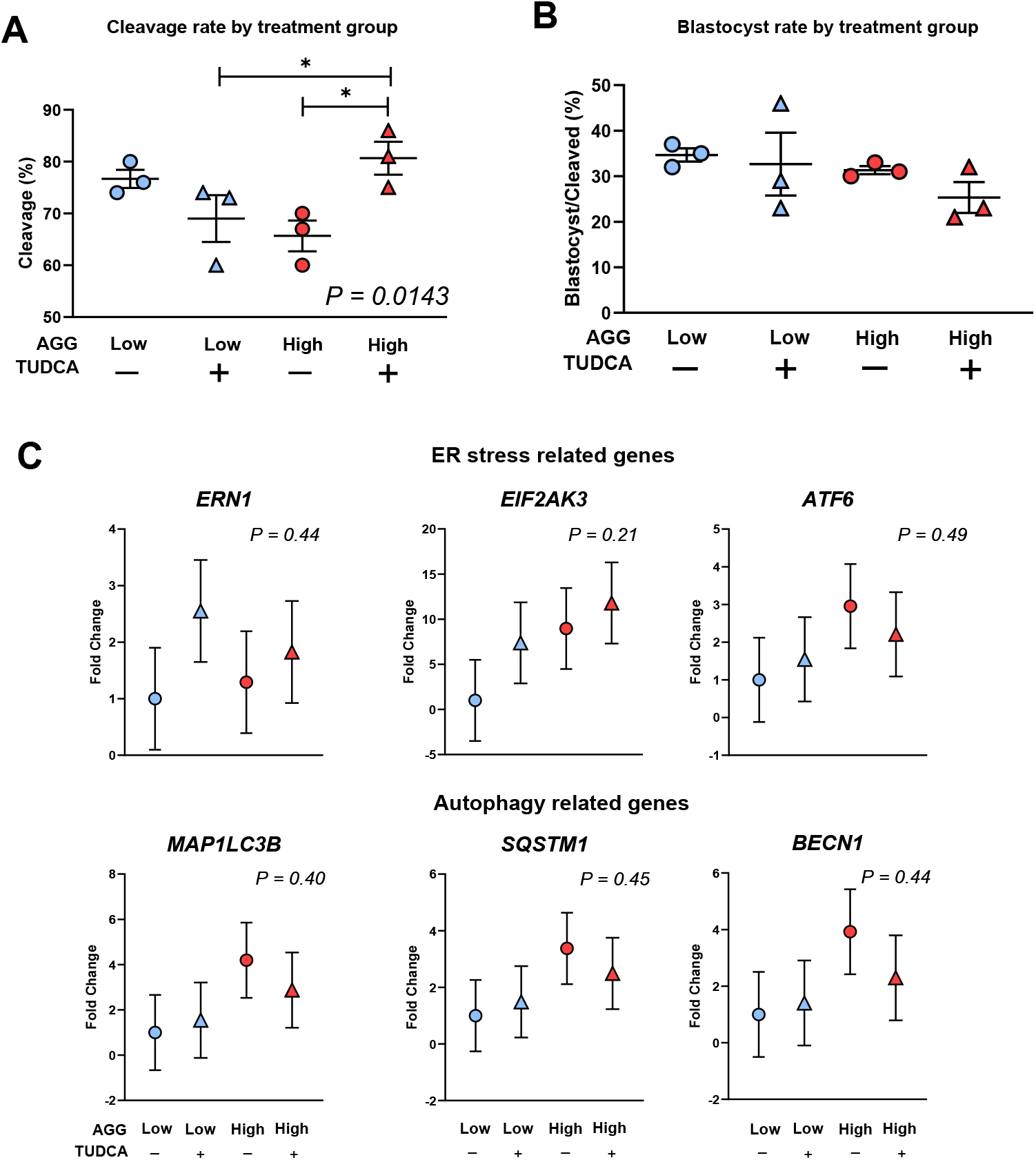
Effect of TUDCA supplementation on embryo development and proteostasis-related gene expression in embryos derived from low- and high-AGG sires. (A) Cleavage rate (%) on Day 3 post-insemination. (B) Blastocyst rate (% of cleaved embryos) on Day 7.5. (C) Relative mRNA expression of ER stress–related genes (ERN1, EIF2AK3, ATF6) and autophagy-related genes (MAP1LC3B, SQSTM1, BECN1) at the 4–6 cell stage, expressed as fold change relative to ACTB (2^™ΔΔCT^, low-AGG –TUDCA as reference). Blue, low-AGG sires; red, high-AGG sires. Bars show mean ± SEM. *P < 0.05.

**Figure S2.**
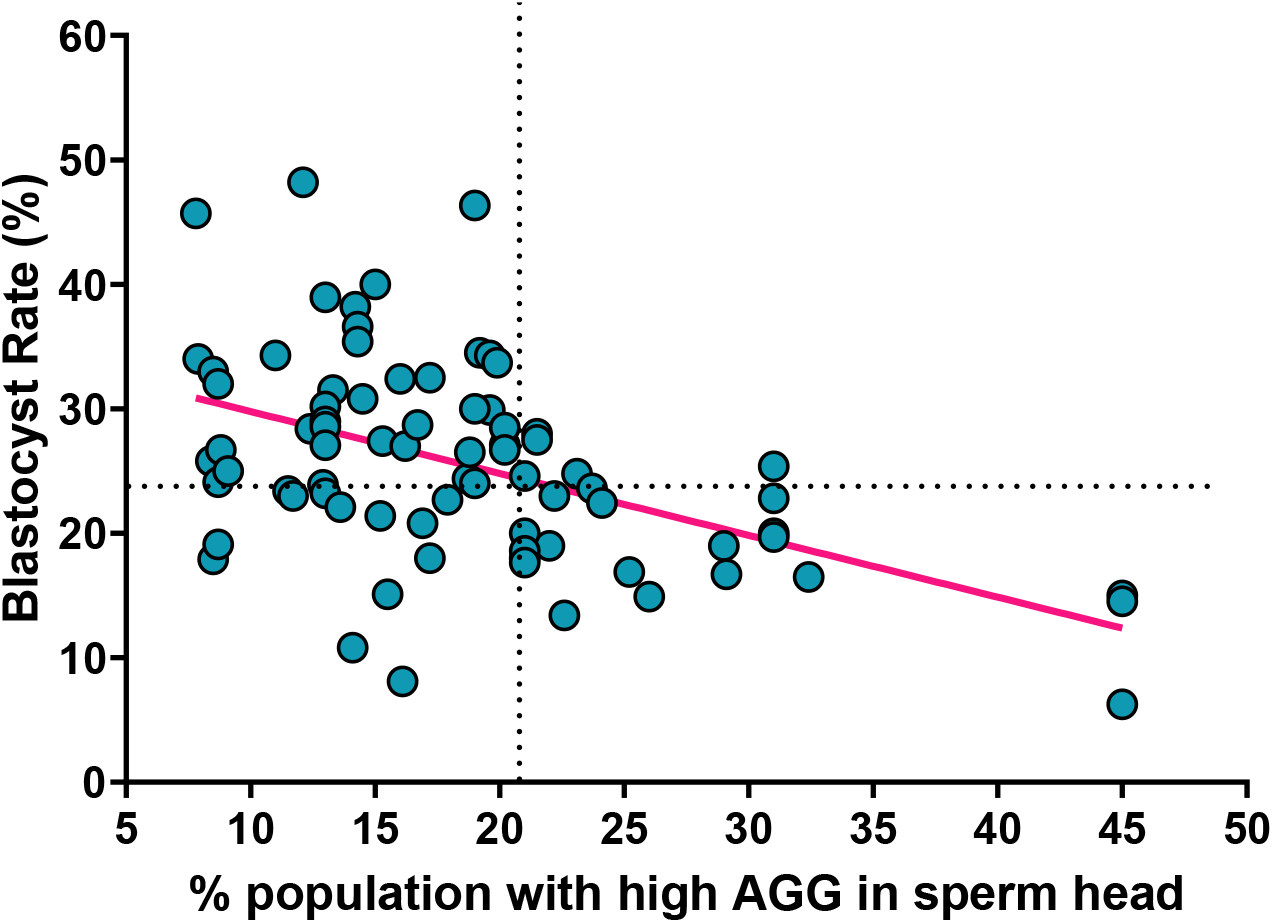
Negative correlation between sperm head AGG content and embryo developmental competence. Blastocyst rate (%) plotted against the percentage of sperm in each ejaculate exhibiting high AGG content in the head. Each point represents an individual sire (n=79).

**Table S1.**
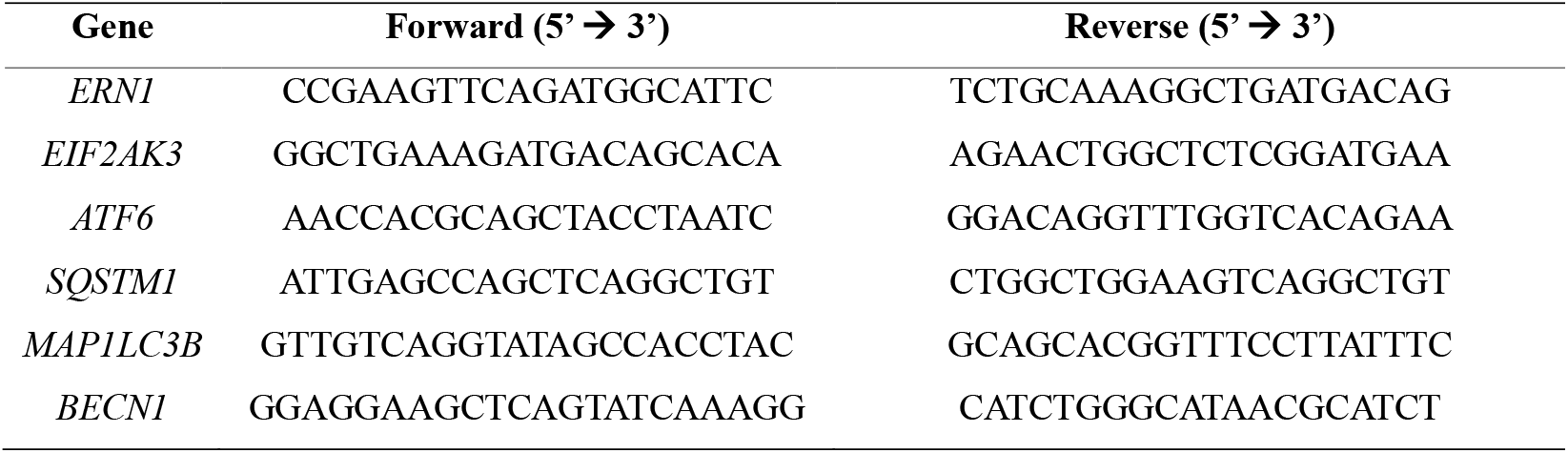
Primer sequences of ER stress and autophagy-related genes.

## Supplemental Material

### Supplemental Video 1

Time-lapse video of embryos produced with either low or high Aggresome content sires. Embryos remained in culture for 8 days.

